# Protein Design Viz (PDV): lightweight protein structure visualization studio with validated quantitative analytics and antibody-specific toolkit

**DOI:** 10.64898/2026.09.08.750039

**Authors:** Konrad Krawczyk, Pawel Dudzic, Sandeep Kumar

## Abstract

**Motivation:** Molecular visualization is dominated by two families of software. Desktop programs such as PyMOL, VMD and UCSF ChimeraX are powerful but heavy to install and operate. Web viewers such as Jmol, 3Dmol.js, NGL, Mol* and iCn3D are light and installation-free, but typically stop at rendering, lacking deeper functionality needed for even basic protein sequence, structural analyses and design tasks. Tools building upon these often lack comprehensive sequence/structure manipulation, surface, interface and antibody-specific analytics functionality or require an external backend. There is a need for a tool that is as frictionless as a web viewer yet carries the analytical depth normally reserved for the desktop or the command line software.

**Results:** We present Protein Design Viz (PDV), a self-contained molecular visualization studio delivered as a single offline HTML file that runs entirely in the browser. Built as an extensive modification of 3Dmol.js, PDV combines a full visualization workflow: multi-object scenes, representations, coloring palette, linked sequence track, publication-quality outline rendering and portable sessions. Additionally we re-implemented four commonly used macromolecular analyses from scratch in client-side JavaScript: a Shrake–Rupley solvent-accessible surface area (SASA) engine, an antibody numbering and germline-assignment engine, a non-covalent interaction detector, and a developability-liability scanner. Each engine is validated against its established reference. PDV’s SASA reproduces FreeSASA at Pearson r ≈ 0.997–0.998 across 2,582 structures spanning proteins, nucleic acids and ligands; its numbering reproduces RIOT for over 99.88% of 1.3 million residue positions across 16,996 sequences and four schemes; its interaction detector reproduces PLIP at macro-F1 0.82, matching Arpeggio as closely as PLIP itself does. PDV brings validated, quantitative structural analysis into a no-install, simple to use tool.

**Availability and implementation:** PDV is a single HTML file, free for noncommercial use under the PolyForm Noncommercial License 1.0.0, available from pdv.naturalantibody.com. It requires only a WebGL-capable browser and runs fully offline.

## 1 Introduction

Three-dimensional visualization of macromolecular structure is a routine step in structural biology, protein engineering and therapeutic antibody discovery. The steady growth of the Protein Data Bank (Berman et al. 2000), together with the plethora of models produced by deep-learning structure predictors (Varadi et al. 2024; Lin et al. 2023), has made interactive inspection of structure a routine activity for a broad user base. The software available to that user base, however, falls into two families, each with its own drawbacks.

The first family is the desktop “heavyweight” such as PyMOL, Visual Molecular Dynamics (VMD), UCSF Chimera and its successor ChimeraX. These pieces of software are mature, powerful and scriptable. They remain the reference tools for publication quality figures and advanced sequence and structural analyses of biological macromolecules (Humphrey et al. 1996; Pettersen et al. 2021). Each of these requires installation and, frequently, a license or registration and carries a substantial memory and disk footprint. They expose much of their analytical functionality through a command language that presents a barrier to occasional users. The boundary between visualization and analysis is, moreover, not sharp: VMD, for example, supports extensive molecular-dynamics trajectory analysis, and integrated commercial platforms such as Schrödinger’s Maestro suite, the Molecular Operating Environment (MOE) and BIOVIA Discovery Studio fold structure visualization into broader modeling, simulation and design environments. Currently, opaque scripting and closed source are also a barrier in developing agentic tools that automate structure visualization workflows. Pieces of software having an open-source component benefit from development of agentic functionality, e.g. extensions/plugins for PyMOL (Golden and Kiefl 2026).

The second family is the web viewer. Jmol (“Jmol: An Open-Source Java Viewer for Chemical Structures in 3D,” n.d.) pioneered in-browser molecular graphics. WebGL-era successors: 3Dmol.js (Rego and Koes 2015), the NGL Viewer (Rose and Hildebrand 2015), Mol* (Sehnal et al. 2021) and iCn3D (Wang et al. 2020) render large assemblies smoothly inside an ordinary browser tab. Web-based servers likewise couple visualization to computation, as in SWISS-MODEL for homology modelling (Waterhouse et al. 2018). The focus here is still on rendering, so it lacks analytical depth. There are pieces of software that build upon such renderers to introduce layers of functionality (Shi et al. 2017; Rego and Koes 2015; Ferla et al. 2020; Tomasello et al. 2020). Nevertheless, user experience (need for server-side communication) and functional depth required for real-world protein-engineering and drug-discovery applications, are often lacking in these applications.

To bridge this gap and to facilitate limited protein analyses via Web-based applications, we have built Protein Design Viz (PDV, pun intended). PDV is a single, self-contained HTML file. It requires neither installation nor network connection beyond the optional convenience of fetching a structure by its PDB identifier. Every structure a user loads is parsed and rendered locally in WebGL, ensuring privacy. Alongside a complete visualization workflow it ships four quantitative analytical engines: (i) solvent-accessible surface area, (ii) antibody numbering, (iii) non-covalent interaction detection, and (iv) in vitro physicochemical degradation liability profiling useful towards developability analyses of therapeutic antibodies and other proteins. Each quantitative module was re-implemented in dependency-free client-side JavaScript so that it runs offline, and validated against the established reference tool in its domain. As such, PDV offers convenience, privacy and analytical depth required to perform the majority of protein design visualization tasks.

## 2 Materials and methods

### 2.1 A single-file application built on WebGL

PDV is distributed as one self-contained HTML document that bundles its entire interface, logic and dependencies. The 3D view is powered by 3Dmol.js (Rego and Koes 2015), the open-source WebGL molecular-rendering library. PDV extensively modifies and extends the basic rendering view with tools to manipulate structure/sequence representation as well as implementation of custom analytical modules such as solvent-accessible surface area (Shrake and Rupley 1973). To ensure that the application runs offline all the dependencies were in-lined in the file or re-implemented in JavaScript, as is the case with antibody numbering, surface accessibility or non-covalent bonding. Structures are loaded by fetching a four-character PDB identifier from the RCSB, by opening local files (PDB, mmCIF, MOL2, SDF or XYZ), or by drag-and-drop; loading is additive and fully multi-object. However, the individual objects can be easily deleted from the session, if and as needed.

Beyond rendering, PDV re-implements four analyses that normally require separate command-line tools, together with exact geometric utilities for distance measurement, proximity-based contact mapping, and structural superposition by the Kabsch algorithm (Kabsch 1976). The four engines, and their reference tools, are: a Shrake–Rupley SASA engine validated against FreeSASA (Mitternacht 2016); an antibody numbering, CDR-delineation and germline-assignment engine that reproduces RIOT (Dudzic et al. 2024); a non-covalent interaction detector modelled on PLIP and Arpeggio (Schake et al. 2025; Jubb et al. 2017); and an antibody physicochemical developability-liability scanner built on Liability Antibody Profiler (LAP) (Satława et al. 2024). Because these engines were re-implemented from scratch in the browser rather than wrapped from existing binaries, their fidelity cannot be assumed. It must be demonstrated via benchmarking against the respective tools as shown in the next section.

Each re-implemented engine was compared against the community-standard tool in its domain, on large and diverse structure sets. The purpose was to establish whether the in-browser implementation reproduces the reference results, rather than to compare computational performance, where the browser versions naturally underperform in comparison to lower-level implementations (e.g. the Rust implementation in RIOT).

### 2.2 Benchmark datasets for the analytical engines

To test the fidelity of our re-implementation, we constructed structure and sequence datasets sampled from the non-redundant antibody, nanobody and protein-complex sets of the NA Structural Database (Chomicz et al. 2026).

The solvent-accessibility and non-covalent-interaction engines were benchmarked on a common structural dataset of 2,582 structures spanning six classes: antibody–antigen, nanobody–antigen, hetero-protein, protein–DNA, protein–RNA and protein–ligand complexes. They comprise 995,373 protein residues and 12,654 non-protein residues (nucleotides, ligands and ions) (Table 1). Using the same set for both engines lets the nucleic-acid and small-molecule classes probe surface accessibility and interface interactions in the same structures. Antibody light chains were de-duplicated so that near-identical variable domains do not dominate the antibody classes. For the non-covalent benchmark the receptor/ligand partition of each class is given in Table 2.

**Table 1.**
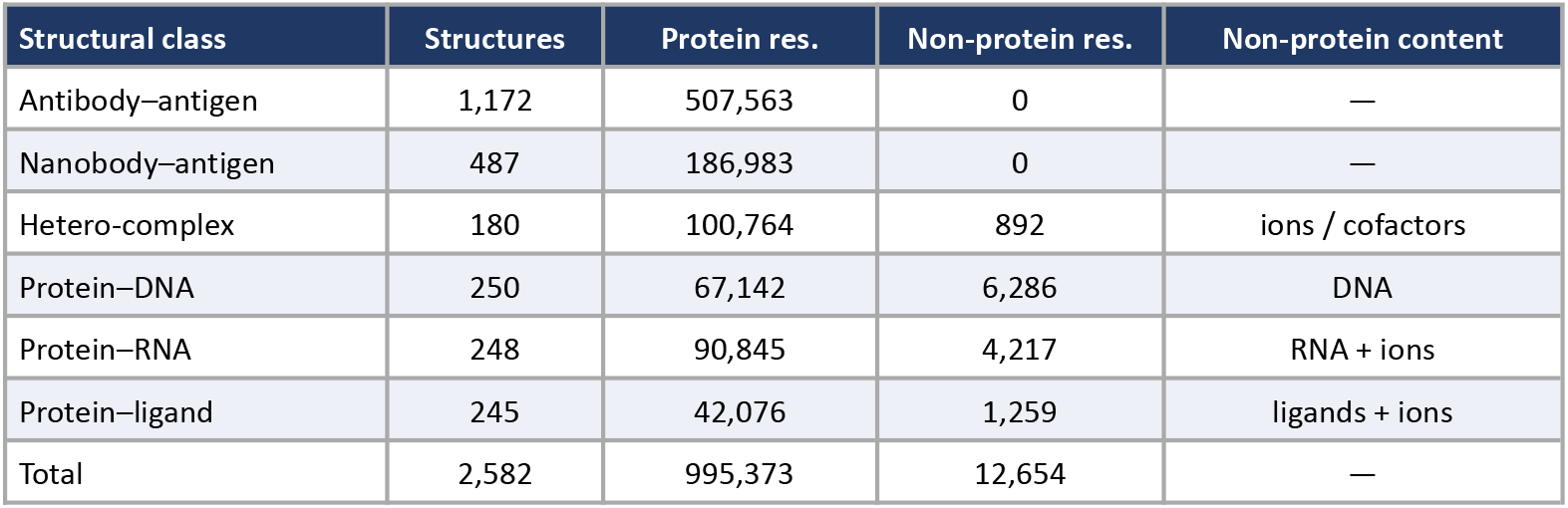
Composition of the shared SASA and non-covalent benchmark set. Residue counts are the per-residue points entering the holo PDV-versus-FreeSASA comparison; non-protein residues are nucleotides, ligands and ions retained in the surface calculation.

| Structural class | Structures | Protein res. | Non-protein res. | Non-protein content |
| --- | --- | --- | --- | --- |
| Antibody–antigen | 1,172 | 507,563 | 0 | — |
| Nanobody–antigen | 487 | 186,983 | 0 | — |
| Hetero-complex | 180 | 100,764 | 892 | ions / cofactors |
| Protein–DNA | 250 | 67,142 | 6,286 | DNA |
| Protein–RNA | 248 | 90,845 | 4,217 | RNA + ions |
| Protein–ligand | 245 | 42,076 | 1,259 | ligands + ions |
| Total | 2,582 | 995,373 | 12,654 | — |

**Table 2.** Receptor/ligand partition used for the non-covalent interaction benchmark, applied to the same structures as Table 1. Protein-interface sets are curated interface files from the NA Structural Database; the ligand, DNA and RNA classes were drawn from the RCSB PDB Search API filtered by resolution and size.

| Class | Receptor / ligand partition (non-covalent benchmark) |
| --- | --- |
| Antibody–antigen | antigen chain = ligand |
| Nanobody–antigen | antigen chain = ligand |
| Protein–protein (hetero) | one chain = ligand |
| Protein–ligand | HETATM small molecule = ligand |
| Protein–DNA | nucleic acid = ligand |
| Protein–RNA | nucleic acid = ligand |

For antibody numbering, 16,996 sequences were used: 2,332 non-redundant antibody complex chains from the NA Structural Database (heavy and light separately) and 487 single-domain camelid antibodies (VHH) as positives, and the 14,177 non-immunoglobulin non-redundant protein monomers available as a specificity negative control.

### 2.3 Solvent-accessible surface area

PDV computes SASA with the Shrake–Rupley numerical algorithm (Shrake and Rupley 1973), implemented as a self-contained Web Worker. Each atom’s van der Waals sphere is expanded by the solvent probe radius (1.4 Å) and sampled with a golden-spiral (Fibonacci) set of 200 test points; a point is buried if it lies inside any neighbouring expanded sphere, and the exposed fraction multiplied by the sphere area gives the atomic accessible area. Neighbour look-ups use a uniform-grid cell list, making the cost linear in atom count. Bondi atomic radii are used (with a 1.5 Å default), hydrogens are excluded, per-atom areas are summed to residues, and relative SASA is obtained by dividing each residue’s area by its theoretical maximum (Tien et al. 2013). A holo/apo toggle computes accessibility either within the whole assembly or for a selection in isolation, and the difference isolates the binding interface. The reimplementation was benchmarked against FreeSASA v2.2.1 (Mitternacht 2016) run in two modes on identical atom sets, the Shrake–Rupley point-sampling algorithm (S–R) and the Lee–Richards slice algorithm (L–R), both with a 1.4 Å probe and the default ProtOr radii. FreeSASA is standard in the field, however its normalization strategy is not perfect as it is based on a fairly old study (Miller et al. 1987). Crystallographic water and hydrogens were removed, while nucleic-acid, ligand and ion atoms were retained so that they contribute to the molecular surface. Agreement was quantified per residue by Pearson and Spearman correlation coefficients, mean and median absolute deviation, RMSD and signed bias (PDV − reference), for both absolute (Å^2^) and relative (0–1) SASA. Non-protein residues, for which relative accessibility is undefined, were compared on absolute SASA only. Residues were labelled buried (relative SASA < 0.20), partially exposed (0.20–0.50) or exposed (≥ 0.50), and label agreement tabulated. For complexes, the chain-ignored change Δ = (apo − holo) relative SASA was compared, and residues with Δ ≥ 0.10 were treated as a binary interface annotation scored by precision, recall, F1 and the Matthews correlation coefficient (MCC).

### 2.4 Antibody numbering

PDV recognises antibody chains, applies a numbering scheme (IMGT, Kabat, Chothia, Martin), delineates the CDRs and assigns germline V and J genes, drawing on the RIOT algorithm (Dudzic et al. 2024). The independent browser implementation was compared against the published RIOT reference across all four supported schemes. For every chain the complete position–residue map was compared, requiring an exact match of both the position label (including insertion codes such as 111.1 or 100A) and the residue at that label; the antibody/non-antibody classification, chain locus and class, species, and germline V/J calls were compared separately, and framework/CDR region boundaries checked per scheme.

### 2.5 Non-covalent interaction detection

PDV’s non-covalent contacts tool classifies each interatomic contact across an interface into a physical interaction type using PLIP-style geometric rules (Salentin et al. 2015). Unlike the reference tools PDV uses no cheminformatics back-end (neither RDKit nor OpenBabel (O’Boyle et al. 2011)). Donor/acceptor and charge assignment come from element identity, residue templates for the standard amino acids and nucleotides, distance-based bond perception, and approximate hydrogen placement. The π/aromatic family is deliberately out of scope, since reliable aromatic-ring perception for arbitrary ligands is not achievable without a full cheminformatics toolkit. The detector was benchmarked against PLIP (Schake et al. 2025; Salentin et al. 2015) and Arpeggio (Jubb et al. 2017). Because the three tools choose different representative atom pairs for the same contact, agreement was scored at the level of the set of binding-site (receptor) residues forming each interaction type, using the residue-level F1, the Dice overlap of the two residue sets, computed pairwise between every pair of tools. Measuring PLIP against Arpeggio on the same structures gives the inter-reference agreement ceiling that bounds any third method.

### 2.6 Developability-liability profiling

PDV’s liability scanner detects common antibody developability chemical degradation motifs: asparagine deamidation, aspartate isomerization, N-glycosylation sequons, methionine and tryptophan oxidation, metal-catalysed histidine oxidation, fragmentation and cleavage sites, and unpaired or missing cysteines. Rather than reporting raw motif counts, each hit is weighted by its structural context. Because chemical degradation is a reaction governed by solvent accessibility and the local structural and sequence environment rather than by evolutionary conservation, solvent exposure is the primary risk determinant: a buried motif is down-weighted because it is inaccessible to solvent. Proximity to the CDRs is used to flag potential impact on binding. The motif catalogue and its context flags follow, to a degree, our Liability Antibody Profiler (LAP) (Satława et al. 2024), which curated roughly 70 severity-graded motifs and, benchmarked against experimental deamidation, isomerization and oxidation datasets, showed that these flags mark about 60% of raw motif hits as low-risk. An exposed, CDR-proximal motif is therefore surfaced as genuinely risky, while a buried motif is down-weighted, and every hit is highlighted jointly on the structure and the sequence track. This engine reuses the two validated engines above rather than introducing new geometry, so its reliability rests on the validations already established.

## 3 Results

### 3.1 Visualization workflow

Every loaded structure (Figure 1) becomes an object with its own visibility, color, name and per-chain controls, and named selections layer representation, coloring and style on top of the base object so that, for example, a whole protein can be shown as cartoon while a binding loop is drawn as ball-and-stick over it. Residues are picked in the 3D scene or on the sequence track, with range and toggle modifiers. Seven representations (cartoon, stick, sphere, ball-and-stick, line, surface and hide) and a broad coloring palette are available. Coloring schemes span spectrum, chain, secondary structure, element, B-factor, charge, several published hydrophobicity scales (Kyte–Doolittle, Eisenberg, Fauchère–Pliska and Wimley–White), residue-type and multiple-sequence-alignment palettes, and solvent accessibility. A continuous sequence track mirrors every coloring scheme and links two-way to the 3D scene through hover, selection and a draggable minimap. These representations apply equally to antibodies, protein complexes, nucleic acids and small-molecule ligands, as illustrated in Figure 2. Publication-quality PNGs are exported with a choice of outline modes, including a line-art and a cel-shaded mode, and up to 4x supersampling. The complete scene, comprising objects, selections, styles and camera position, can be written to a portable session file.

**Figure 1.**
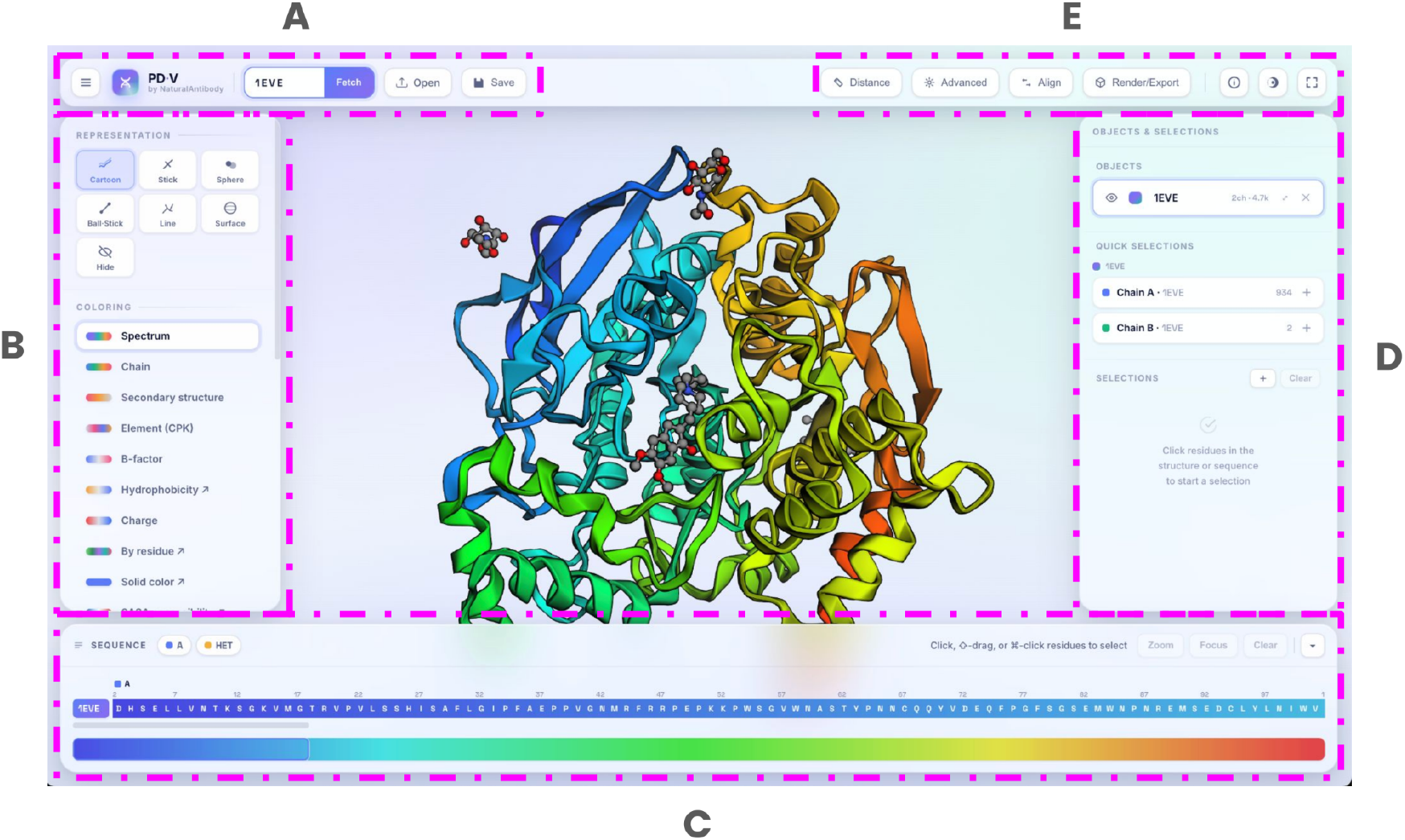
The PDV workspace. A) Structure bar - loading structures and saving the current state. B) Representation and coloring, allowing for changing between color schemes (solid color, SASA, hydrophobicity, charge per-residue schemes, etc.) and representation (cartoon, ball& stick etc.). C) Sequence track to facilitate selections in 1D. D) Objects & selections, to manipulate different subsets of a given objects or sets of objects. E) Tools - miscellaneous tools: distance measurement, alignment, advanced analytics and publication-grade rendering.

**Figure 2.**
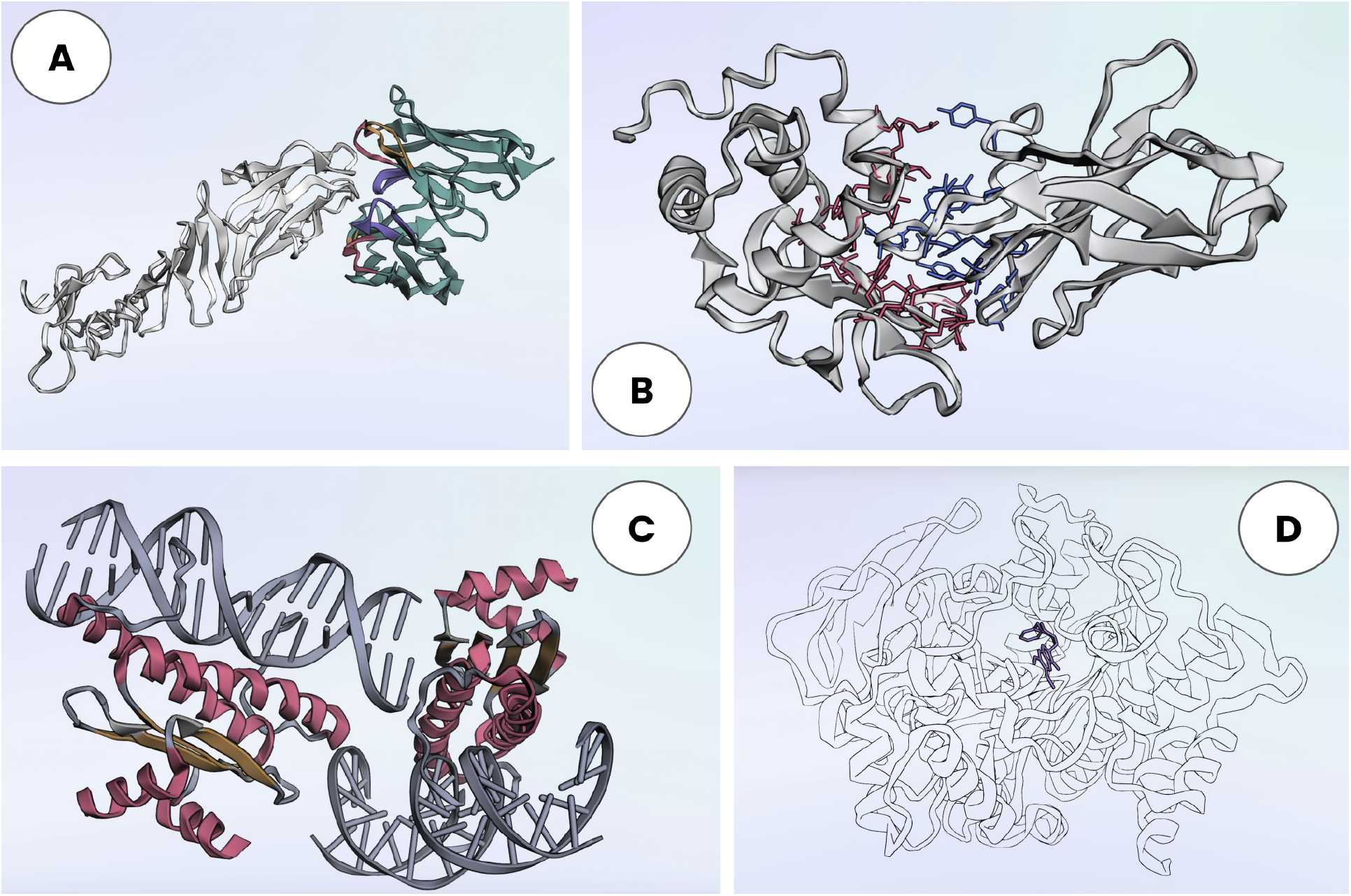
Versatility of PDV molecular representations. A) Antibody CDR and framework delineation. B) Identification of contacts between protein chains. C) Visualization of nucleic acids. D) Visualization of small molecule ligands.

### 3.2 Solvent-accessible surface area versus FreeSASA

We tested 2,582 structures spanning six classes, protein–DNA, protein–RNA and protein–ligand, protein-protein, protein-antibody and protein-nanobody complexes (see Table 1) on the agreement between our JavaScript re-implementation of surface accessibility and FreeSASA. Both pieces of software agree almost perfectly on protein per-residue accessibility (Figure 3). For absolute SASA in the holo state the Pearson correlation was 0.997 against the Shrake–Rupley reference and 0.994–0.998 against Lee–Richards, uniformly across all six structural classes (Table 3). Spearman correlations were essentially identical (ρ ≈ 0.99 throughout), confirming that the agreement is not an artefact of a few high-exposure residues. Switching FreeSASA from Shrake–Rupley to Lee–Richards changes the correlation only in the third decimal place, confirming that the small residual differences originate from the radius set rather than the surface-sampling scheme. PDV is marginally more expansive than FreeSASA, with a small constant positive bias of 0.40–0.76 Å^2^ per residue depending on class, consistent with the slightly larger Bondi radii relative to ProtOr. Computed under a normalization (Tien et al. 2013) with maximum-ASA table, capped at 1.0, relative SASA performs matches well FreeSASA implementation (Pearson 0.996–0.997; mean absolute difference 0.013–0.014 on the 0–1 scale); the larger offset seen when the two engines are compared in their native relative conventions (off diagonals in Figure 3) is because FreeSASA does not cap the ratio at 1.0 and it differs in the maximal values used for the normalization, using (Miller et al. 1987) .

**Figure 3.**
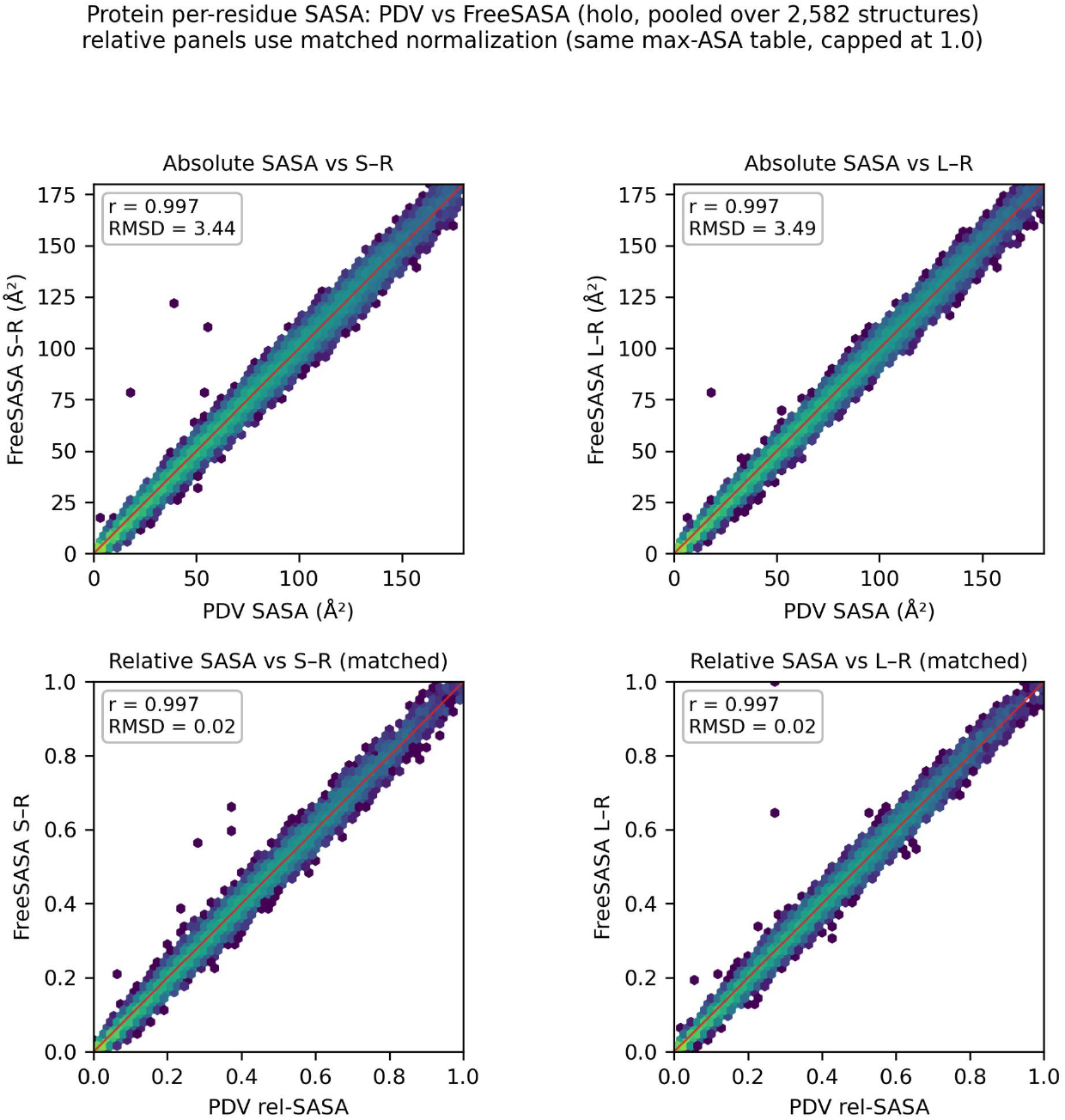
Protein per-residue SASA agreement (holo, pooled over all 2,582 structures). Log-density hexbins; red line is identity. Top: absolute SASA against FreeSASA Shrake–Rupley and Lee–Richards. Bottom: relative SASA under matched normalization (both engines normalised to the Tien 2013 max-ASA table and capped at 1.0). PDV is tightly collinear with both FreeSASA algorithms. The few off-diagonal points arise from a handful of residues (≈0.003%), modelled in alternate conformations, where PDV includes all deposited conformers (which mutually occlude and lower the area) while FreeSASA evaluates a single one, and cases where FreeSASA’s per-residue reporting attributes a bound ligand’s or nucleotide’s area to a protein residue that shares its residue number on another chain.

**Table 3.**
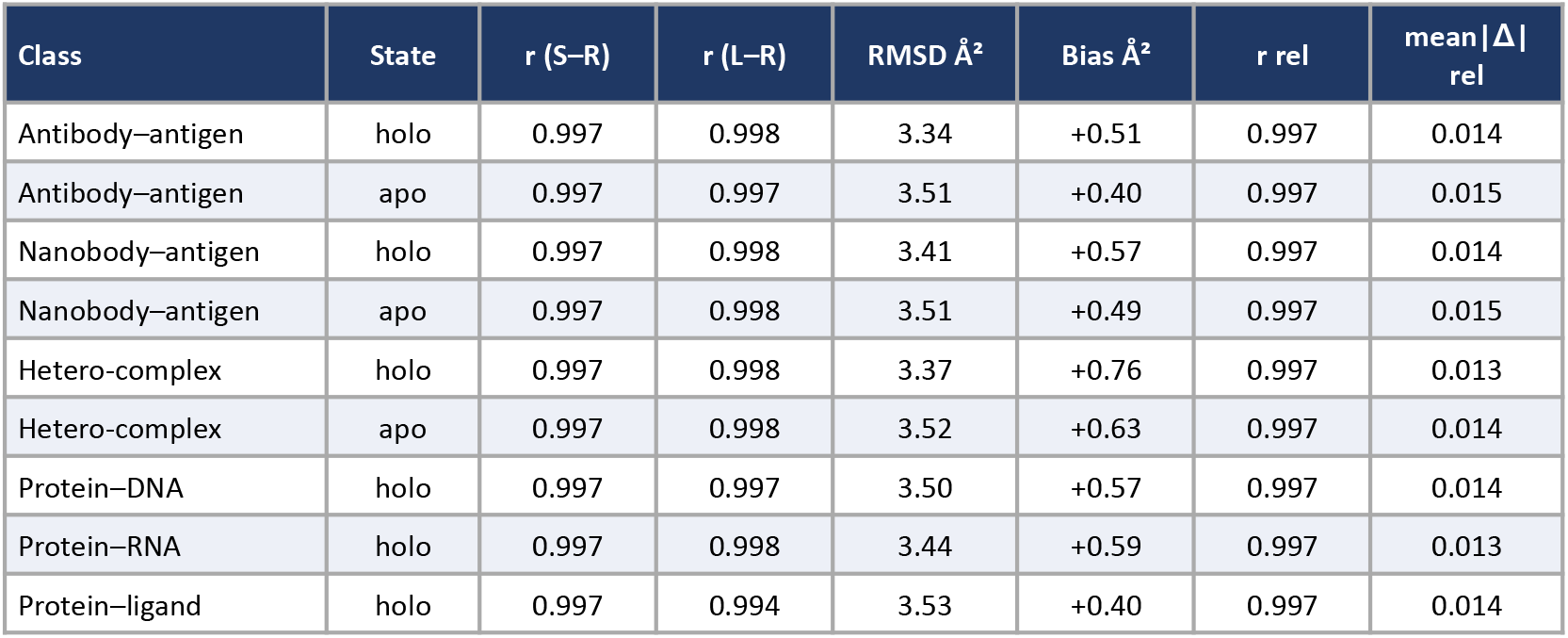
Protein per-residue SASA agreement between PDV and FreeSASA across the 2,582-structure benchmark (representative states shown; apo reported for antibody, nanobody and hetero classes). RMSD and bias are for absolute SASA (Å^2^; bias = PDV − FreeSASA, Shrake–Rupley); r rel and mean|Δ| rel are for relative SASA under matched normalization. S–R, Shrake–Rupley; L–R, Lee–Richards.

The equivalence extends to the non-protein atoms. On nucleotide, ligand and ion residues the two engines correlate at r = 0.994–0.999 (Table 4, Figure 4), with a larger positive bias of 1.79–3.53 Å^2^ per residue because PDV and FreeSASA assign systematically different radii to phosphate, sugar and exotic ligand atoms; the expected radius-set effect rather than an algorithmic disagreement, which does not propagate to the neighbouring protein residues. Reduced to the three-state buried/partial/exposed labelling used for downstream annotation, PDV and FreeSASA assign the same label to 89.4–91.3% of protein residues, with the DNA-, RNA- and ligand-bound categories indistinguishable from the protein-only ones and disagreements concentrated at the 0.20 and 0.50 thresholds where a sub-Ångström difference in area flips a borderline residue between adjacent classes.

**Table 4.** Absolute SASA agreement on non-protein residues (nucleotides, ligands, ions) in the holo state. The larger bias than for protein residues reflects the different radii the two engines assign to these atoms; the correlation remains ≥ 0.99 throughout.

| Class | n residues | $r$ (S–R) | $r$ (L–R) | RMSD $\text{\AA}^2$ | Bias $\text{\AA}^2$ |
| --- | --- | --- | --- | --- | --- |
| Protein–DNA | 6,286 | 0.995 | 0.994 | 8.17 | +2.58 |
| Protein–RNA | 4,217 | 0.994 | 0.994 | 8.72 | +3.53 |
| Protein–ligand | 1,259 | 0.998 | 0.992 | 5.55 | +1.79 |
| Hetero-complex | 892 | 0.999 | 1.000 | 5.38 | +2.44 |

**Figure 4.**
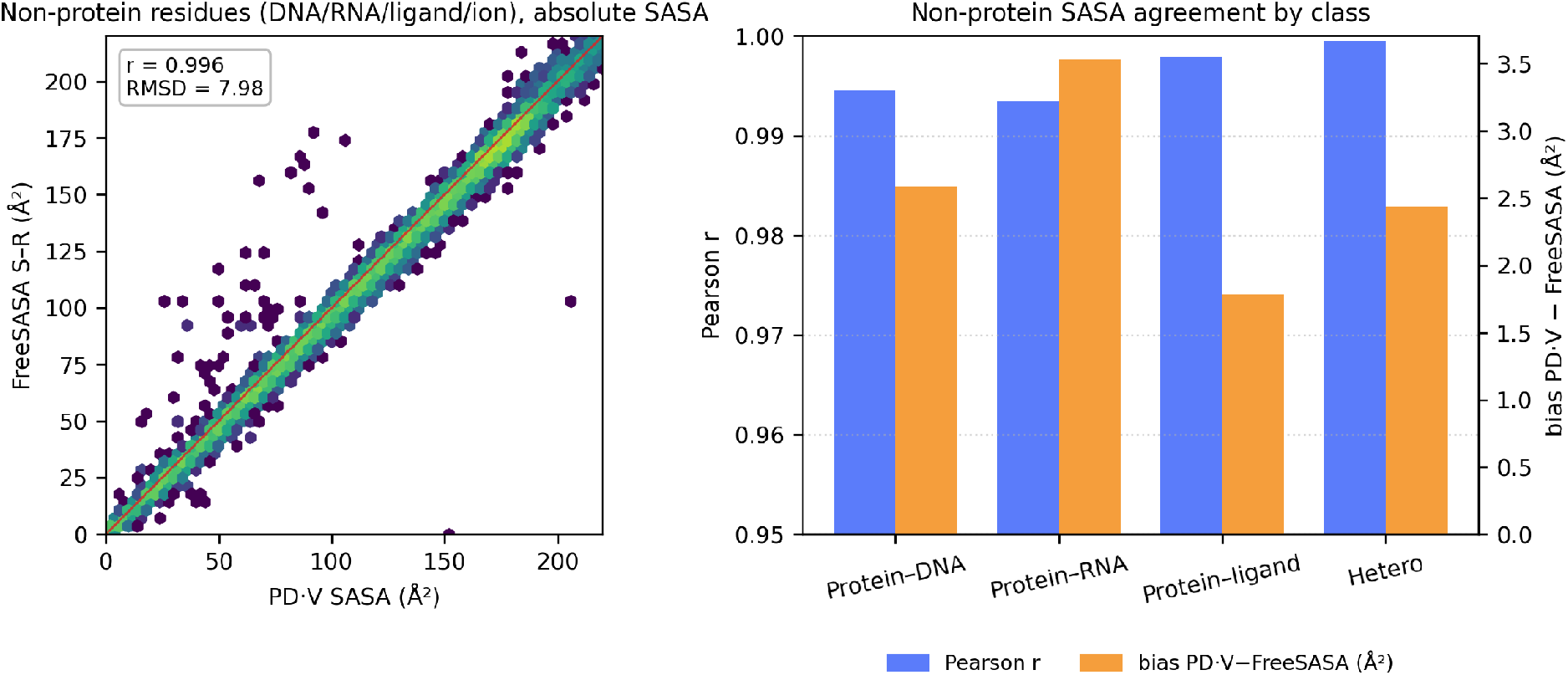
Non-protein absolute SASA. Left: PDV versus FreeSASA (S–R), pooled over DNA/RNA/ligand/ion residues. Right: per-class Pearson r (left axis) and mean bias (right axis); correlation stays ≥ 0.99 while the constant offset grows for phosphate-rich nucleotides.

A central use of the SASA is to flag interface residues by the accessibility a chain loses when its partners are present (Esmaielbeiki et al. 2016). Removing partner chains and recomputing accessibility, PDV and FreeSASA reproduce this signal almost identically: the per-residue Δ(apo − holo) relative SASA correlates at r ≥ 0.988 for every class, and treating residues with Δ ≥ 0.10 as a binary interface annotation, PDV recovers the FreeSASA-defined interface at F1 0.954–0.968 and MCC ≥ 0.95, with high precision (Table 5, Figure 5). This holds for protein–DNA, protein–RNA and protein–ligand interfaces as well as protein–protein ones, confirming that the two engines agree on how a nucleic-acid or ligand partner occludes the protein surface. We therefore treat PDV’s SASA values as interchangeable with FreeSASA for downstream analysis, accessible via the representation coloring panel (Figure 6).

**Table 5.** Agreement between PDV and FreeSASA on the interface annotation obtained by removing partner chains, across all six classes. Δ r, Pearson correlation of per-residue Δ(apo − holo) relative SASA; the remaining columns treat residues with Δ ≥ 0.10 as interface and score PDV against the FreeSASA labelling.

| Class | $\Delta r$ (S–R) | Agreement | Precision | Recall | F1 | MCC |
| --- | --- | --- | --- | --- | --- | --- |
| Antibody–antigen | 0.996 | 0.992 | 0.997 | 0.942 | 0.968 | 0.964 |
| Nanobody–antigen | 0.997 | 0.994 | 0.996 | 0.924 | 0.958 | 0.956 |
| Hetero-complex | 0.996 | 0.993 | 0.997 | 0.937 | 0.966 | 0.962 |
| Protein–DNA | 0.995 | 0.991 | 0.996 | 0.932 | 0.963 | 0.958 |
| Protein–RNA | 0.996 | 0.991 | 0.994 | 0.933 | 0.963 | 0.958 |
| Protein–ligand | 0.988 | 0.993 | 0.992 | 0.919 | 0.954 | 0.951 |

**Figure 5.**
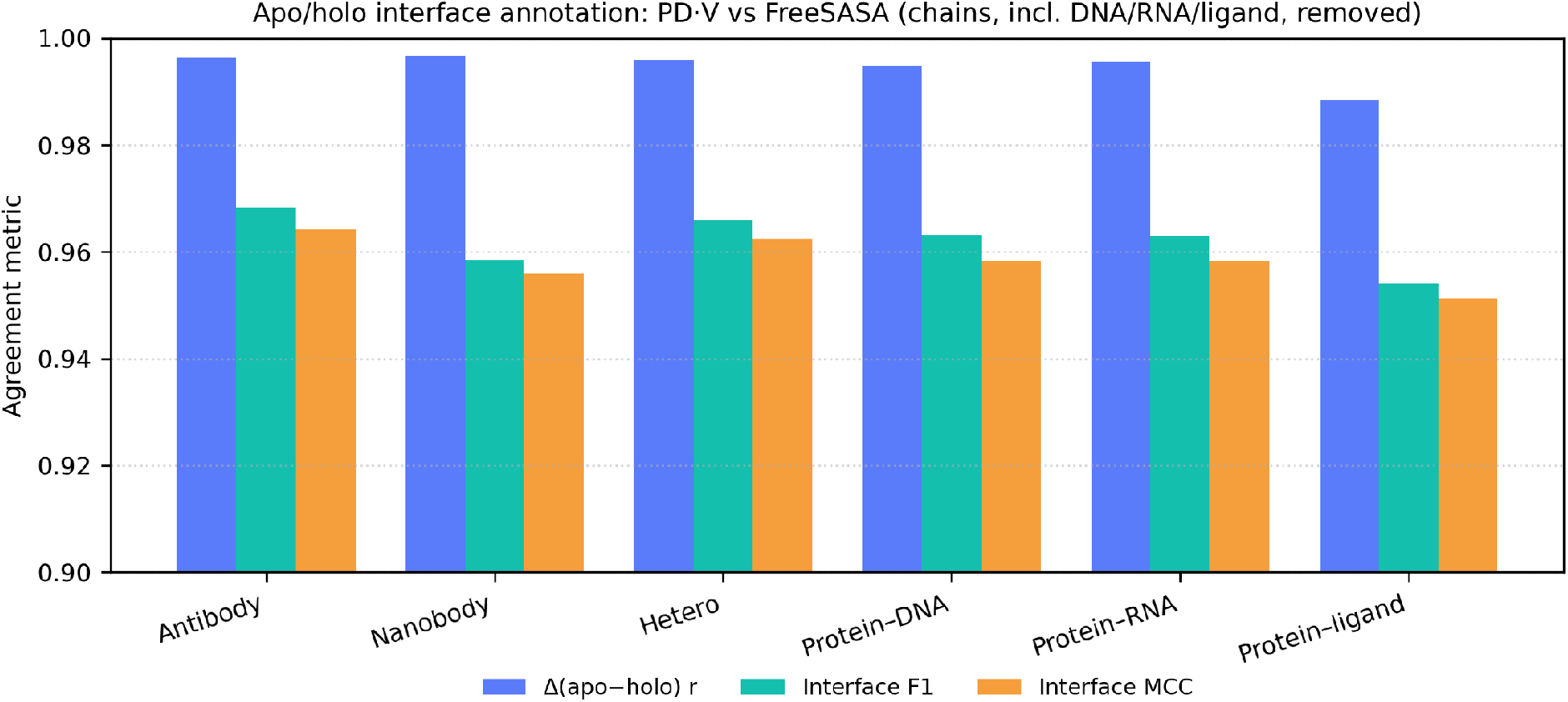
Apo/holo interface reproduction across the six classes. Correlation of the per-residue accessibility change (blue) and agreement of the binary interface annotation (F1, MCC). Protein–DNA, protein–RNA and protein–ligand interfaces are recovered as accurately as protein–protein ones.

**Figure 6.**
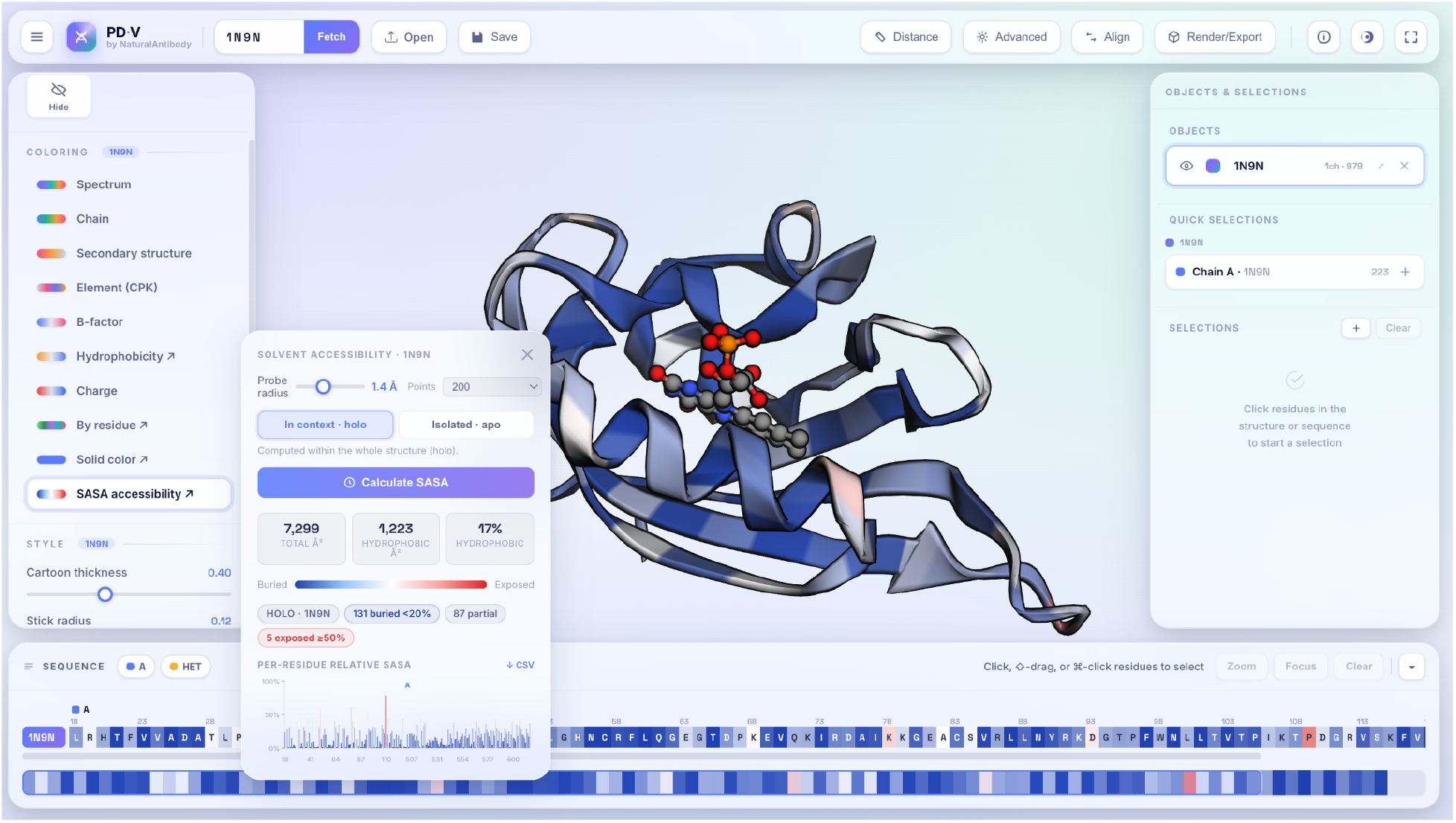
The SASA panel. Probe radius, sampling density and a holo/apo context toggle drive the Shrake–Rupley engine; readouts include total and hydrophobic area, buried/partial/exposed counts, a per-residue bar chart and CSV export. Structure and sequence are colored blue→white→red by relative exposure.

### 3.3 Antibody numbering versus RIOT

The antibody/non-antibody classification matched RIOT for all 16,996 sequences in the benchmark dataset; the two classes are separated by the V-gene alignment significance (Figure 7). For the 2,819 antibody and nanobody chains, chain locus and heavy/light class matched RIOT for 100% of chains, species for 99.29%, the V germline gene for 97.69% and the J germline gene for 98.12% (Figure 8). Requiring an exact match of both the position label (including insertion codes) and the residue at that label, per-position agreement exceeded 99.88% under every scheme, and 98.4–98.7% of chains were numbered identically to RIOT at every single position (Table 6). Agreement on the error-prone insertion-coded positions ranged from 98.3% to 99.4%, and PDV reproduces IMGT-specific reverse-ordered insertions at positions 33, 61 and 112. Restricting attention to chains where both tools selected the same germlines raised per-position agreement to 99.93-–99.94%, confirming that the small residual differences arise from germline tie-breaks on heavily mutated sequences rather than from the numbering procedure.

**Figure 7.**
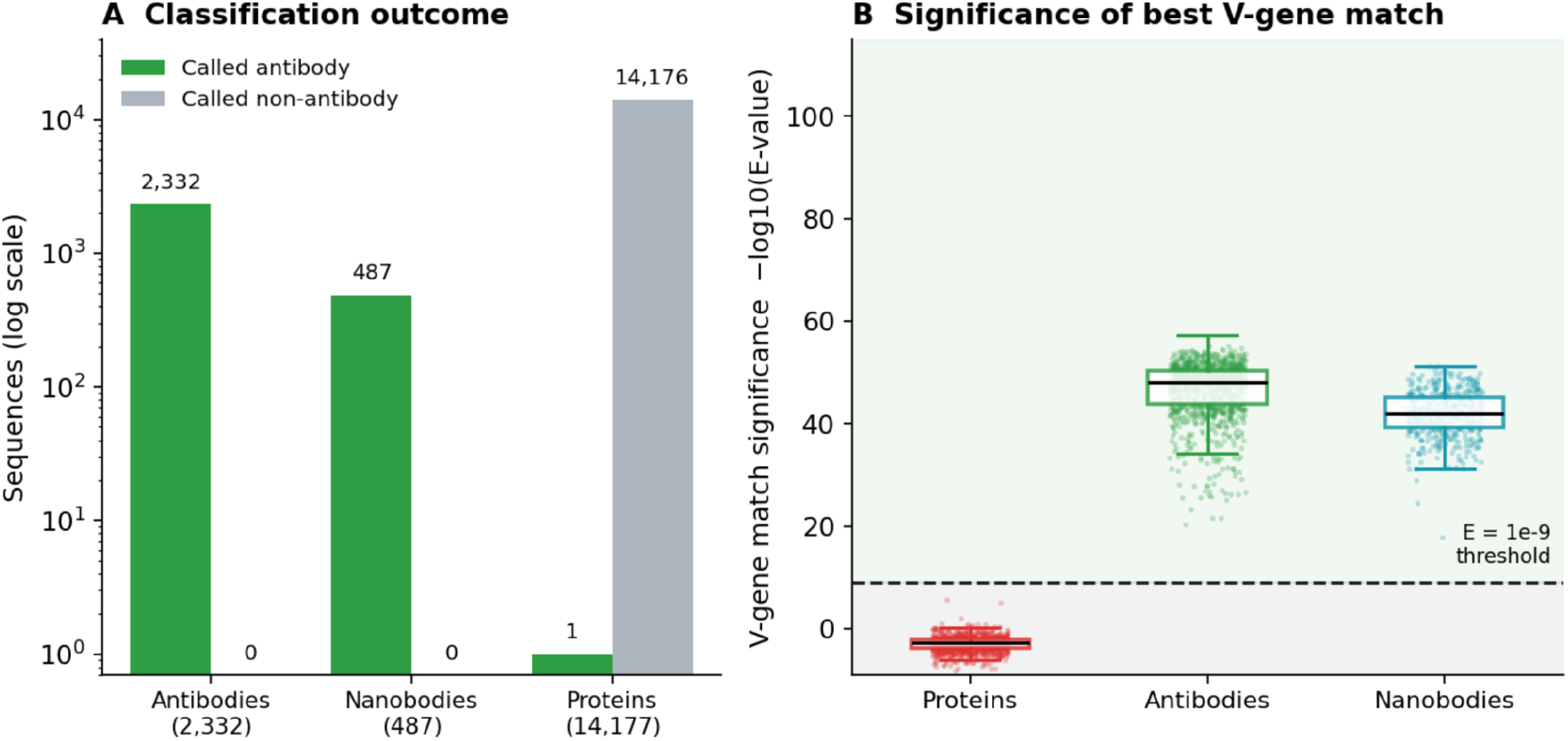
Antibody detection. Antibody detection by V-gene alignment significance. (A) Classification outcome (log scale): all 2,332 antibodies and 487 nanobodies are called antibodies, and 14,176 of 14,177 non-antibody proteins are correctly rejected. The single protein called an antibody (PDB 6RUL) carries a genuine immunoglobulin VHH domain. (B) Significance of the best V-gene match per sequence, -log10(E-value); the dashed line is the E = 1e-9 decision threshold, separating the ‘antibody’ zone (green, above) from the ‘non-antibody’ zone (grey, below). Antibodies and nanobodies fall far inside the antibody zone (medians 48 and 42), while every protein with a measurable alignment sits well inside the non-antibody zone (median -2.8); a further 10,221 of the 14,177 proteins produced no V-gene alignment at all and lie off-scale below. Boxes show median and interquartile range; points are individual sequences (proteins subsampled for display). The lone protein above the threshold is the 6RUL Ig-domain protein.

**Table 6.** Residue-numbering concordance between PDV and RIOT per scheme (2,819 antibody and nanobody chains). Per-position: exact label-plus-residue matches. Identical chains: chains matching RIOT at every position. Insertion codes: agreement restricted to inserted positions. Region boundaries: chains with all seven framework/CDR boundaries identical.

| Scheme | Per-position | Identical chains | Insertion codes | Region boundaries |
| --- | --- | --- | --- | --- |
| IMGT | 99.897% | 98.37% | 98.29% | 98.86% |
| Kabat | 99.888% | 98.55% | 99.17% | 99.40% |
| Chothia | 99.906% | 98.65% | 99.40% | 99.18% |
| Martin | 99.905% | 98.65% | 99.35% | 99.18% |

**Figure 8.**
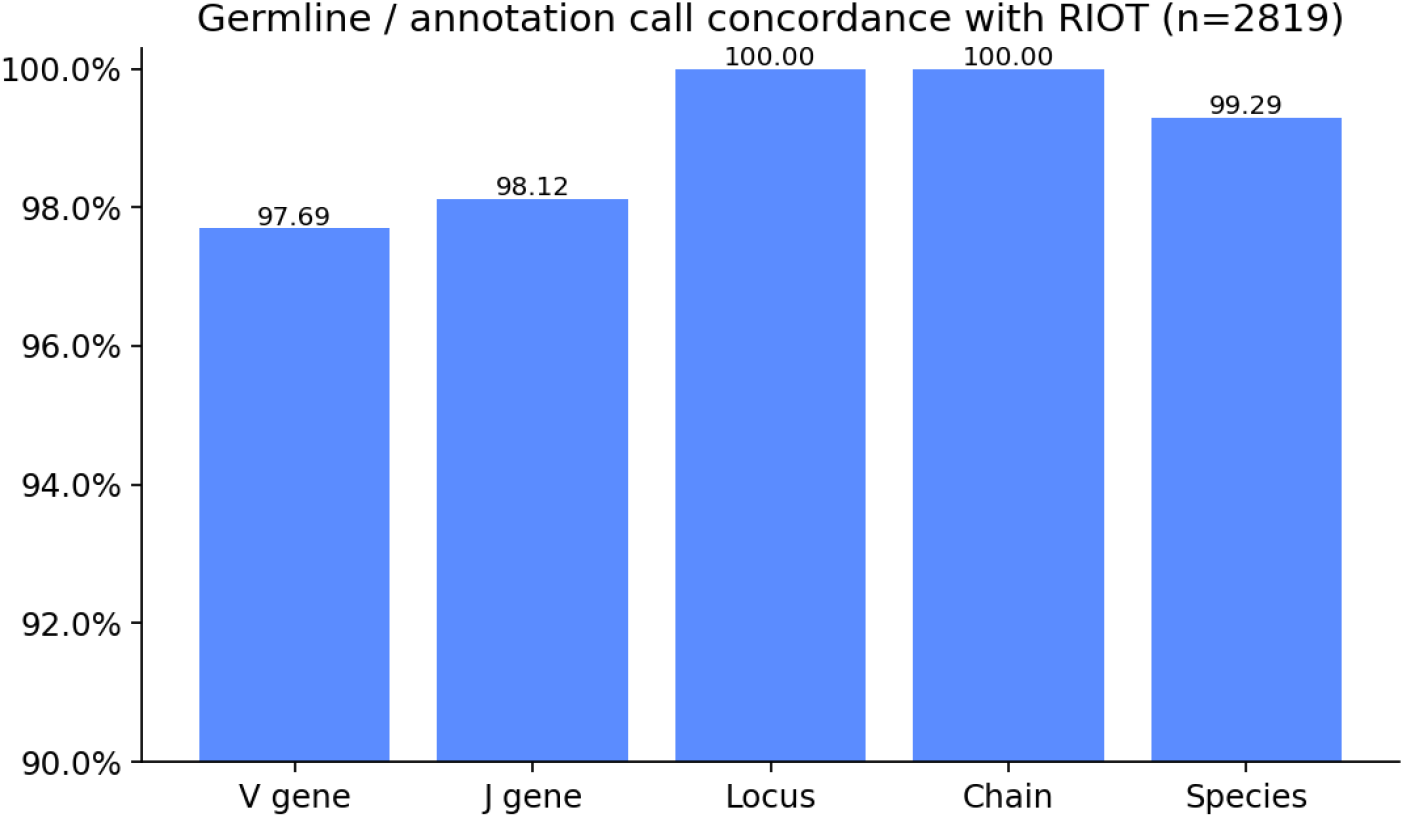
Germline and annotation concordance with RIOT. Concordance of germline (V, J) and chain-annotation (locus, chain class, species) calls across 2,819 antibody and nanobody chains. The residual V/J differences are choices between closely homologous germlines that leave the numbering essentially unaffected.

### 3.4 Non-covalent interactions versus PLIP and Arpeggio

PDV covers the non-covalent interactions at the interfaces that are a subset of those detected by Arpeggio and PLIP. The coverage of all three tools is compared in Table 7. Unlike PLIP and Arpeggio, PDV omits the π/aromatic family, which requires aromatic-ring perception. PLIP and Arpeggio (both using OpenBabel) agree with each other only at F1 0.61–0.69 overall on their shared interaction types, and as low as 0.12–0.27 on small-molecule ligands. PDV reproduces PLIP at macro-F1 0.82, reaching F1 0.93 on hydrophobic contacts, 0.81 on hydrogen bonds and 0.89 on salt bridges, and 0.78–0.81 on halogen bonds and metal complexes from element identity and geometry alone (Table 8, Figure 9). It agrees with Arpeggio at essentially the same level the two references agree with one another, and slightly higher on hydrophobic and hydrogen interactions. Resolved by structural class, agreement is high on protein and protein–protein interfaces dominated by the twenty standard residues PDV templates directly (F1 0.79–0.92 against PLIP) and lower on small-molecule ligands and nucleic acids, where chemical perception is indispensable and where the two references themselves diverge most. The typed interactions are therefore best treated as a fast, well-calibrated guide rather than definitive assignments, especially for small-molecule ligands. The non-covalent bonds as well as interaction distance can be visualized in PDV on the structure as well as using the Sankey diagram for convenience (Figure 10).

**Table 7.** Interaction-type coverage across the three tools (✓ covered; — not covered). PDV covers the geometry-defined core without a cheminformatics back-end and omits the π/aromatic family. PDV additionally annotates van der Waals contacts and disulfides for visualization; these were not part of the quantitative benchmark.

| Interaction type | Detection | PDV | PLIP | Arpeggio |
| --- | --- | --- | --- | --- |
| Hydrophobic | Close approach of two apolar carbons | ✓ | ✓ | ✓ |
| Hydrogen bonds | Donor and acceptor must be in proximity; The angle of donor, hydrogen and acceptor (DHA) must be >120° | ✓ | ✓ | ✓ |
| Weak Hydrogen bonds (C–H...O) | Weak donors (C–H) with relaxed distance and angle thresholds. | ✓ | — | ✓ |
| Ionic / salt bridges | Close proximity of groups with opposite charge; salt bridges additionally share the hydrogen bond. | ✓ | ✓ | ✓ |
| Halogen bonds | C–X...acceptor σ-hole (X = F/Cl/Br/I) | ✓ | ✓ | ✓ |
| Metal complexes | Metal ion coordinated by O/N/S donors | ✓ | ✓ | — |
| Water bridges | Donor and acceptor linked through a water | ✓ | ✓ | — |
| π-stacking / π-cation | Aromatic ring–ring or cation–aromatic | — | ✓ | ✓ |
| Extended $\pi$ (C-/donor-/S-π, amide-ring) | Weak contacts to aromatic systems | — | — | ✓ |

**Table 8.** Pairwise residue-level F1 among PDV, PLIP and Arpeggio, pooled over all classes. Dashes mark interaction types a tool does not report. The PLIP–Arpeggio column is the inter-reference ceiling: PDV matches PLIP well above that ceiling and tracks Arpeggio about as closely as PLIP itself does.

| Interaction type | PDV–PLIP | PDV–Arpeggio | PLIP–Arpeggio | Counts (PLIP/Arp) |
| --- | --- | --- | --- | --- |
| Hydrophobic | 0.93 | 0.74 | 0.69 | 11.6k / 17.7k |
| Hydrogen bonds | 0.81 | 0.66 | 0.61 | 24.4k / 24.0k |
| Weak H-bonds (C–H...O) | — | 0.52 | — | — / 19.6k |
| Salt bridges | 0.89 | 0.67 | 0.69 | 6.9k / 4.2k |
| Halogen bonds | 0.78 | 0.47 | 0.65 | 27 / 13 |
| Metal complexes | 0.81 | — | — | 3.2k / — |
| Water bridges | 0.68 | — | — | 9.5k / — |

**Figure 9.**
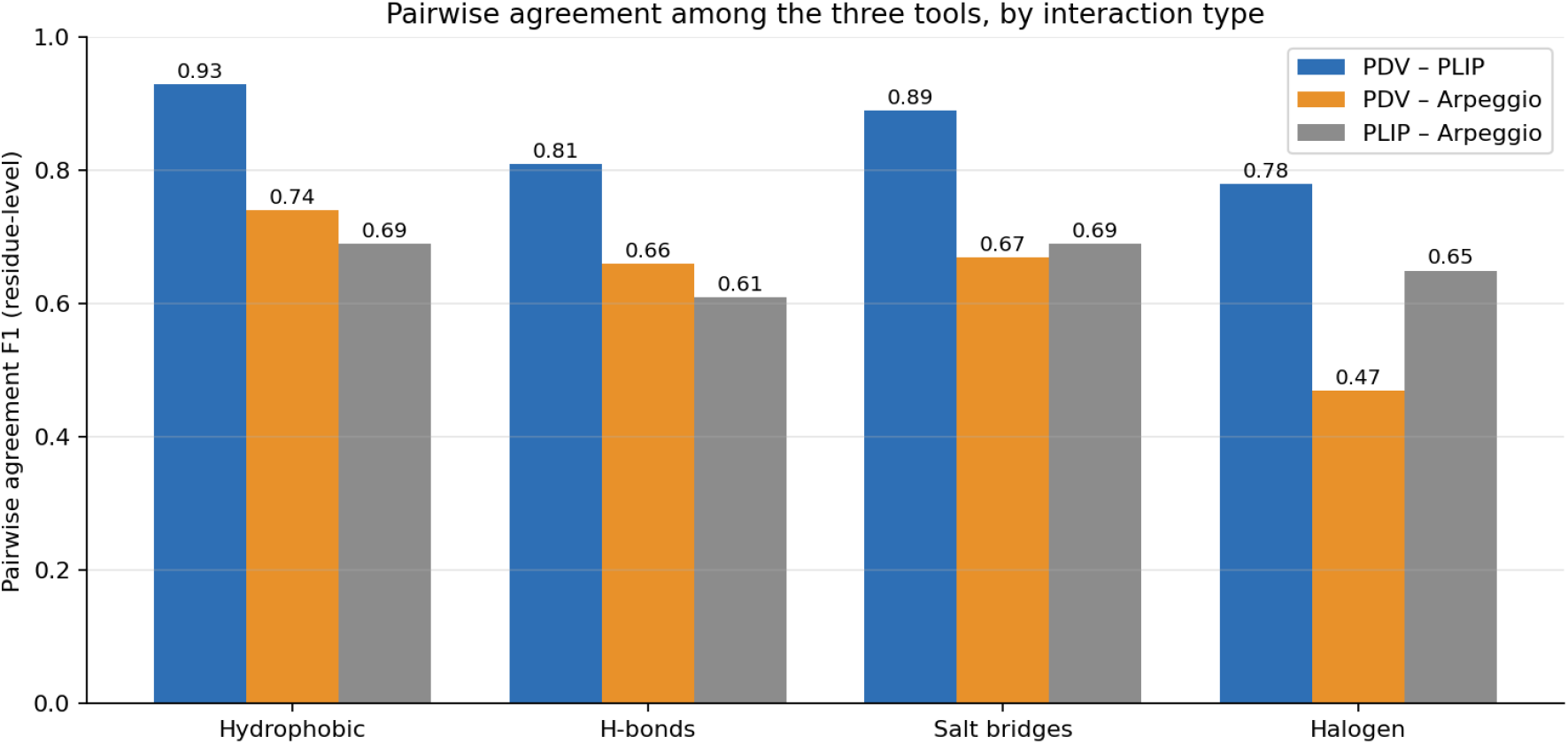
Pairwise agreement by interaction type. Residue-level F1 among the three tools for the four shared interaction types. PDV’s agreement with PLIP (blue) sits well above the PLIP–Arpeggio inter-reference ceiling (grey), while PDV–Arpeggio (orange) matches, and for hydrophobic and hydrogen bonds exceeds, that ceiling.

**Figure 10.**
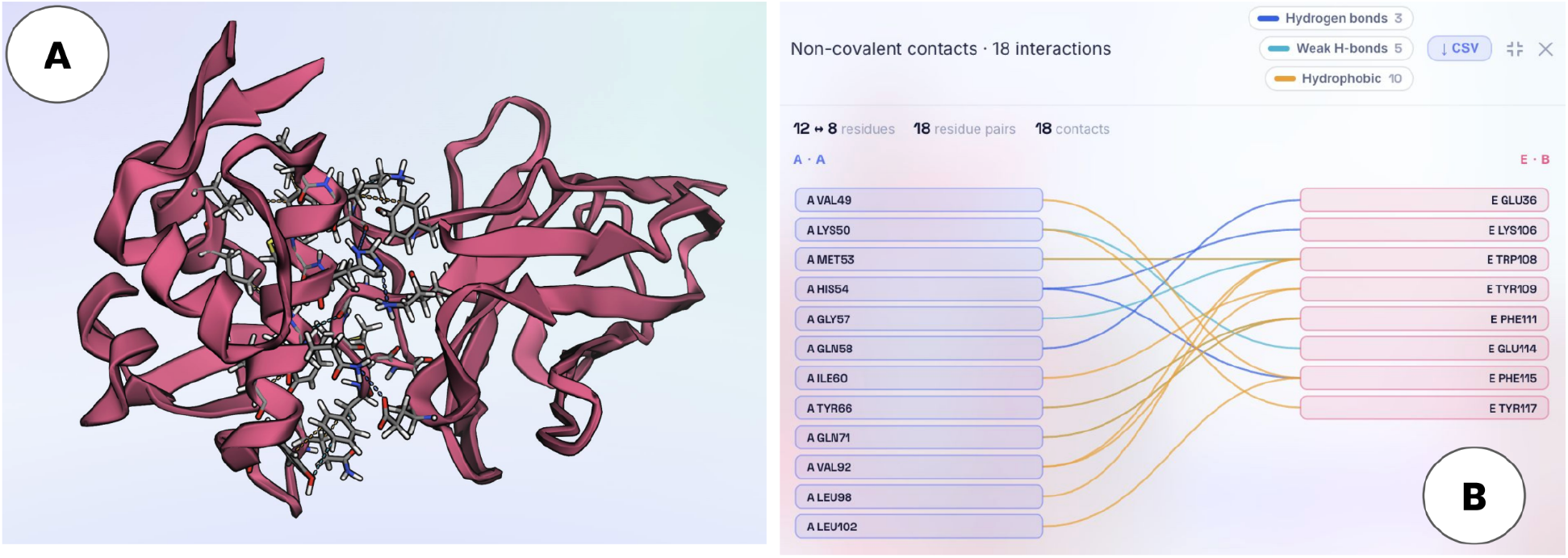
The non-covalent contact map. A) Detected non-covalent contacts are visualized in the structure. B) Residue pairs across two groups are rendered as a Sankey diagram colored by interaction type rather than by distance; the interacting residues are saved as selections and exported to CSV.

## 4 Discussion and conclusion

By developing PDV we show that one can achieve computational convenience without sacrificing analytical accuracy as demonstrated by high concordance between the four analytical engines as implemented within PDV and their respective versions running outside of the web viewer.

Our method is intrinsically limited by the absence of a cheminformatics back-end for small-molecule interaction typing and the π/aromatic family. Nonetheless, we do not envisage this as a fundamental hindrance, as the software can be further developed pursuing deeper analytics for small molecules, nucleic acids and the like, when handled by individuals versed in the field. We further note lack of Molecular Dynamics trajectory support, which of course can be added in future versions.

As it stands, PDV makes validated, quantitative structural analysis easily available, and we hope it is useful to the protein-design and antibody-engineering communities in the current form.

